# Reduced confidence intervals and novel candidate genes for quantitative trait loci associated with apple scab resistance in *Malus domestica*

**DOI:** 10.64898/2026.04.08.717319

**Authors:** Romane Lapous, Camille Haquet, Caroline Denancé, Juliette Bénéjam, Laure Perchepied, Kaat Hellyn, Hélène Muranty, Charles-Eric Durel, Julie Ferreira de Carvalho

## Abstract

Apple scab, caused by *Venturia inaequalis*, remains one of the most damaging diseases in apple orchards, driving intensive pesticide use worldwide. Reducing this dependence requires the deployment of durable resistance, ideally through the combination of major resistance genes (R genes) with quantitative trait loci (QTL) that confer partial and potentially complementary protection. Yet, few apple scab QTLs have been functionally validated, and their underlying mechanisms remain largely unresolved.

Here, we refined and functionally described, with transcriptomic data, five resistance QTLs in a biparental population of 1,970 individuals derived from the cross ‘TN 10-8’ × ‘Fiesta’. Using 43 newly developed KASP markers, QTL locations were substantially precised through high-resolution genotyping and phenotyping with two *V. inaequalis* isolates exhibiting contrasting virulence. Four QTL (qT1, qF11, qF17, qT13) were validated, while qF3 was not confirmed. Transcriptomic data comparison revealed the expression of candidate genes within the narrowed intervals, including receptor-like proteins in qT1, and RNAi- and signaling-related genes in qF11 and qF17, suggesting a diversified and complementary defense network.

These findings refine the genetic architecture of apple scab resistance and suppose the existence of shared molecular pathways between major R gene, such as the well-described *Rvi6* gene, and quantitative resistance, with for instance the QTL qT1. The identified loci and markers provide robust tools for marker-assisted and genomic breeding aimed at developing apple cultivars with complementary and potentially durable resistance pathways.

## Introduction

*Venturia inaequalis*, the causal agent of apple scab, remains the most economically damaging fungal disease of apple worldwide, causing substantial yield losses and requiring intensive fungicide applications in temperate regions ^1^. The pathogen overwinters in fallen leaves, producing ascospores that initiate primary infections on young leaves and fruits during spring. Subsequent cycles of conidial dissemination lead to secondary infections, resulting in defoliation, fruit deformation, and, in severe cases, tree weakening ^2^. To limit the environmental and health impacts of repeated fungicide use, the development of genetically resistant cultivars has become an essential strategy for sustainable apple production ^3^.

Historically, breeding for scab resistance has relied on monogenic resistance conferred by resistance genes or R genes. Up to now, at least 18 such loci (*Rvi1*–*Rvi18*) have been identified from cultivated and wild *Malus* accessions ^4–7^. The first and most widely used, *Rvi6* (previously named *Vf*), originates from *Malus floribunda* and encodes a leucine-rich repeat receptor-like protein (LRR-RLP) ^8–10^. Approximately 90% of resistant cultivars released since the 1980s carry this *Rvi6* locus ^11^. However, some *V. inaequalis* populations are able to overcome this resistance, as firstly reported in the cultivar ‘Prima’ in Germany in 1988 ^12^. The origin of such populations could result from the migration of virulent isolates from non-European apples, and their maintenance in environmental disease reservoirs, like ornamental crabapples ^13^. This breakdown underscores the limitations of monogenic resistance, which imposes strong selection pressure on pathogen populations.

In contrast, quantitative resistance, controlled by quantitative trait loci (QTL), typically provides partial but presumably more durable protection, covering functions that might complement R genes ^14^. Each locus contributes modestly to resistance, and the combined effect of multiple QTLs may slow pathogen adaptation and enhance resistance durability if they mobilize different mechanisms. To date, at least 14 apple scab QTLs have been reported ^7^, yet only a few have been precisely mapped or functionally characterized due to broad confidence intervals (CIs) and limited marker density in earlier studies. Among these, three QTL—qT1, qF11, and qF17—have been repeatedly detected and partially characterized in biparental populations derived from the cultivars ‘TN10-8’ and ‘Fiesta’ ^15–18^. The locus qT1, located on chromosome 1 and derived from ‘TN10-8’, co-localizes with *Rvi6*, which has been shown to be a member of an extracellular leucine-rich repeat receptor family ^8,9^. In contrast, qF11 and qF17, both originating from ‘Fiesta’, reside on chromosomes 11 and 17, respectively, and exhibit a strong synergistic interaction, reducing disease severity only when both resistance alleles are combined ^18,19^. Interestingly, these two QTLs were effective against a broader range of isolates than qT1 ^16,18^. Moreover, the pyramiding of qT1, qF11, and qF17 in a ‘TN10-8 × Fiesta’ progeny has been shown to block *V. inaequalis* development at multiple infection stages—from penetration to sporulation ^20^—although subsequent pathogen adaptation has eroded the protection conferred by qF11 and qF17 ^19^. More recently, two additional QTLs—qT13 on chromosome 13 (from ‘TN10-8’) and qF3 on chromosome 3 (from ‘Fiesta’)—were reported by Bénéjam et al. (2021)^18^ but remained poorly resolved due to wide CIs and low SNP density.

To elucidate their genetic basis and exploit them for breeding, it is essential to increase mapping precision and identify putative causal genes. Fine-mapping approaches rely on larger offspring populations (∼500 to >10,000 individuals when available), thereby increasing the number of recombination events ^21^. Coupled with SNP genotyping—the most common source of DNA sequence variation—these approaches help reduce QTL CIs, provide new markers for marker-assisted selection (MAS), limit the co-introgression of undesirable genes, and decrease the number of candidate genes for functional validation ^22–25^. In addition to these approaches, integrating ‘-omics’ data—such as genomics, transcriptomics, proteomics, and metabolomics—can help prioritize candidate genes for functional validation. Such integration may involve sequence alignment ^21^, gene expression analyses ^25^, or the identification of QTL co-localizations ^26–28^.

In this study, we analyzed an extended ‘TN10-8 × Fiesta’ progeny of 1,970 individuals to refine the genetic localization of five QTL (qT1, qF11, qF17, qT13, and qF3) involved in quantitative resistance to *V. inaequalis*. Using newly developed KASP markers and phenotyping with two *V. inaequalis* isolates of contrasting virulence, we improved the mapping resolution of the QTLs. Then, the integration of transcriptomic data from Bénéjam et al. (2024) ^29^ enable us to propose candidate genes potentially involved in apple scab resistance. These findings improve our understanding of the genetic basis of quantitative resistance in apple and provide molecular resources for marker-assisted and genomics-assisted breeding strategies aimed at combining complementary resistance mechanisms for durable scab resistance.

## Materials and methods

### Plant material

Seeds of the cross between apple genotype ‘TN10-8’ and cultivar ‘Fiesta’ (referred to as the ‘TxF’ progeny) were produced in the framework of the DARE European project (Durable Apple Resistance in Europe, 1998-2002). A part of these seeds were firstly elevated in greenhouse until their implantation in orchards, then allowing their phenotyping through grafting to decipher the genetic control of scab resistance segregating in this material ^16^. This initial population, hereafter referred as base population, is composed of 267 individuals and allowed for the detection of three QTLs of ‘Fiesta’ and two of ‘TN10-8’ ^15,16,18,30,31^. ‘Fiesta’ is an offspring from a cross between the cultivars ‘Cox’s Orange Pippin’ (resistant to apple scab) and ‘Idared’ (susceptible to apple scab) and is carrying resistance and susceptibility alleles on each of chromosomes 3, 11 and 17 that are subsequently referred as qF3, qF11 and qF17, respectively. The hybrid ‘TN10-8’, is coming from a cross between the French heirloom cultivar ‘Reinette Clochard’ and a descendent from the Russian cultivar ‘Antonovka’ (fairly resistant to apple scab) ^32^. This genotype is carrying resistance and susceptibility alleles on each of chromosome 1 (qT1) and 13 (qT13). The MUNQ (Malus UNiQue genotype code), as described by Muranty et al. (2020)^33^ and Durel et al. (2023)^34^, was 2396 for ‘TN10-8’, 763 for ‘Fiesta, 163 for ‘Cox’s Orange Pippin’, 717 for ‘Idared’, 118 for ‘Reinette Clochard’ and 4 for Antonovka.

The remaining seeds from the ‘TxF’ progeny, stored at −20°C, were used in the present study and each seed still corresponds to a unique genotype. This population, hereafter referred as extended population, is composed of 1,970 seeds. In the fall 2022, seeds were germinated, transplanted into soil, and subsequently cultivated in a greenhouse on their own roots. Young leaf discs of each individual were sampled using the “BioArkTM Leaf collection kit” from LGC Biosearch Technologies^TM^ company (United Kingdom, UK) for subsequent genotyping.

### Identification of QTL boundaries, development of SNP markers and qPCR-based assays

In previous studies, QTL locations were assessed using 267 individuals ^18^, genotyped with SNPs from the 20K SNP arrays ^35,36^. Context sequences of 43 SNPs localized upstream, within, and downstream of the five QTL CIs, and heterozygous in ‘Fiesta’ or ‘TN10-8’, were retrieved from both the 20K and 480K SNP arrays and blasted against the reference genome of ‘Golden Delicious’ doubled haploid GDDH13 ^37^ to then be visualized using JBrowse ^38^. This step allowed the assignment of broader context sequence surrounding each candidate SNP (mean length 150 bp; range 105-216 bp). Additional SNPs, identified by aligning the sequencing data used to develop the 480K array ^36^ to the GDDH13 genome ^37^, were also considered in order to develop adapted primers for KASP genotyping.

Leaf material was sent to LGC Biosearch Technologies^TM^ (Herts, UK) for DNA extraction and SNP genotyping using KASP technology. For each selected SNP, two allele specific forward primers and one common reverse primer were designed and validated by LGC Biosearch Technologies^TM^ (Herts, UK). The assay mix preparation and PCR amplifications were performed according to the user’s guide and manual (LGC Biosearch Technologies^TM^, Hoddesdon, Herts, UK). The concentration of DNA samples was measured using NanoDrop2000 spectrophotometer (Thermo Fisher Scientific, Waltham, MA, USA). DNA samples were diluted for genotyping, as required. KASP genotyping was carried out on a StepOnePlusTM real-time PCR system (Thermo Fisher Scientific, Waltham, MA, USA).

Genotyping data were cleaned and analyzed using SNPviewer™ (LGC Biosearch Technologies, UK) at INRAE (Angers), which visualizes KASP results as cluster plots. Marker quality was evaluated based on clustering performance and genotype discrimination among seedlings derived from two genotyped parents. Out of the 43 markers tested, 36 were validated. The seven non validated markers exhibited poor clustering, likely resulting from suboptimal primer hybridization due to imperfect primer design or additional polymorphisms near the target SNP.

Using the same DNA extracts, additional SNP markers, were developed at INRAE to further densify the CIs and peak of four QTLs (qT1, qF11, qF17, and qT13). A second set of seven SNP markers was successfully designed and incorporated. KASP primers were designed at Eurofins Genomics. These markers were analyzed on the ANAN genotyping platform (Univ. Angers, France). Genotyping was then done using the PACE® mix (3crbio, Harlow, UK) following manufacturer protocol. Thus, a total of 43 SNPs distributed along the five QTLs of interest were used. Each SNP is presented with sequence context, SNP of interest position and additional SNPs in S1 File. This file also contains virtual markers, added with computational analyses described below.

### Apple scab phenotypic scoring and analyses

#### *Venturia inaequalis* isolate inoculum and apple scab scoring

The seedlings described above were own-rooted and non-replicated; therefore, scab inoculations were performed sequentially on the same plants, which were pruned between each inoculation. Seedlings were randomly distributed into four blocks in the greenhouse.

Two *V. inaequalis* isolates were used: the reference isolate ‘EU-B04’ (Origin: Belgium, host: ‘Golden Delicious’) previously described in Caffier et al. (2015) ^39^ and Le Cam et al. (2019) ^40^ and the isolate ‘09BCZ014’ (Origin: France, host: ‘TN10-8’ × ‘Prima’ progeny), referred to as isolate ‘2557’ in Laloi et al. (2017) ^20^. The isolate ‘09BCZ014’ partially overcomes the resistance conferred by the qT1 resistance QTL segregating in the ‘TxF’ progeny. The use of this isolate allows to better study the effect of other QTLs segregating in the progeny.

Monoconidial suspensions were prepared from diseased dry leaves at a concentration of 2.5 × 10^5^ conidia.ml^−1^ and sprayed on seedlings. Inoculated plants were then incubated for two days at 17°C under a plastic sheet to maintain high humidity, according to the conditions described by Caffier et al. (2010) ^41^. The percentage of leaf surface exhibiting sporulating lesions was scored at 14 and 21 days post-inoculation (dpi) using the ordinal scale (0 to 7) described in Calenge et al. (2004) ^16^. Three experiments were conducted from January to May 2022 (Table 1). One experiment was performed with isolate ‘EU-B04’ (coded ‘Vi-B04’) for which additional scoring of resistance symptoms (leaf chlorosis and crispation) was recorded at 14 dpi using the same ordinal scale (0 to 7 scores, reflecting the percentage of foliar surface covered with chlorosis or crisped). Two other experiments were then conducted using the isolate ‘09BCZ014’ (coded ‘Vi-Z14_1’ and ‘Vi-Z14_2’). ‘Vi-Z14_1’ was performed on all the ‘TxF’ progeny seedlings (1,970) whereas ‘Vi-Z14_2’ was restricted to individuals recombining in CIs of QTLs of interest (1,101). For ‘Vi-Z14’ experiments, only leaf sporulation was recorded (susceptibility symptoms).

**Table 1.** Summary of experiment characteristics.

|  | Experiment 1 | Experiment 2 | Experiment 3 |
| --- | --- | --- | --- |
| **Name** | Vi-B04 | Vi-Z14_1 | Vi-Z14_2 |
| **Month** | January | March | May |
| **Seedling number** | 1,970 | 1,970 | 1,101 |
| **Vi isolate** | EU-B04 | 09BCZ014 | 09BCZ014 |
| **Phenotyping** | Sporulation at 14<br>and 21 dpi<br>Chlorosis at 14 dpi<br>Crispation at 14 dpi | Sporulation at 14<br>and 21 dpi | Sporulation at 14<br>and 21 dpi |

Dpi, days post-inoculation.

Using sporulation scores at both dates *i.e* 14 dpi and 21 dpi, the Area Under Disease Progression Curve (AUDPC) was calculated as a quantitative summary of disease severity over the course of the infection using the following equation:

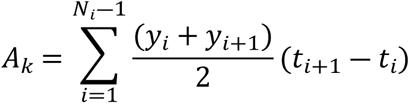

Where *y_i_* is the disease score at the *i*th day of observation and *t* is the number of days after *V. inaequalis* inoculation at the *i*th observation. For AUDPC calculation, the initial time point (t_0_) was set to 0.

All statistical analyses were performed using the software R V2022.07.2+576 (R Core Team, 2022). ANOVAs (function *aov* from the *stats* package) were conducted to test for assessor and block effects and, when necessary, were followed by Tuckey post hoc tests (function *tukey_hsd* from the *rstatix* package). Although each block consisted of genetically distinct individuals, their mean genetic value was assumed to be equivalent due to the random distribution of individuals among blocks, allowing adjustment of phenotypic variables for block, assessor, and experiment effects, with the experiment effect included only for the Vi-Z14 experiments. For the ‘Vi-B04’ experiment, QTL mapping was performed using the adjusted AUDPC, chlorosis and crispation scores (hereafter ‘AUDPC_B04’, ‘CHLOROSIS_B04’ and ‘CRISPATION_B04’). For the ‘Vi-Z14’ experiments, adjusted AUDPC additionally accounted for variations between the two experiments (‘AUDPC Vi-Z14_adj’). Phenotyping data used for QTL mapping are available in S2 File.

Distributions and correlations between studied variables were examined with ggplot function (*ggplot2* package) and ggpairs function (*GGally* package), respectively. In the ‘Vi-B04’ experiment, the strong effect of the major QTL qT1, detected on LG1 of ‘TN10-8’ with the reference isolate ‘EU-B04’ ^16,18^, is hiding the more moderate effects of other QTLs. Therefore, the ‘TxF’ progeny was further subdivided into two subsets of individuals according to the presence or absence of the resistance allele at qT1 predicted by SNP data at the QTL peak. AUDPC distributions were then observed on two distinct subpopulations: one including the entire extended progeny (1,970 individuals) and a second, named ‘qT1-’ including 721 individuals carrying the susceptible allele at qT1.

### Mapping and genetic analyses of the targeted QTLs

#### Genetic maps

Genotyping data were analyzed using SNPviewer software provided by LGC Biosearch Technologies^TM^ with manual correction whenever necessary to improve dataset quality. The order of SNP physical positions was checked against the latest apple reference genetic map ^42^. A linkage map was constructed for each parent with JoinMap 4.1 software ^43^. In total, over the five targeted regions, 43 SNP markers were used and the associated genetic map is available in S3 File.

#### QTL mapping

QTL analyses were conducted using the R/qtl package ^44^. In addition to SNP markers, ‘virtual’ markers were added with *calc.genoprob* and *sim.geno* functions using 1,000 simulations in order to have one marker per centiMorgan (cM). Simple interval mapping and composite interval mapping were then performed using a multiple imputation method and a normal distribution model (using *cim* function). The logarithm of the odd (LOD) score threshold to identify the statistically significant QTLs was determined using 1,000 permutations (α = 0.05). LOD thresholds were approximately 1.80 for all variables.

The LOD score, the 2-LOD support CI, and the contribution of each QTL to the overall phenotypic variance (individual R^2^) were extracted from R/qtl analyses, together with the global QTL contribution (global R^2^). Individual and global R² were calculated with the *fitqtl* function (used for fitting a defined multiple-QTL model). Interactions between QTLs were studied by variance analysis using the genotyping data of each SNP closest to the peak of each QTL, and were detailed by the *effectplot* function. These results were used to define the model for the calculation of the global R^2^ with the *fitqtl* function. Graphics with LOD score and positions of markers were constructed using MapChart 2.32 ^45^.

### Identification of candidate genes

Sequence contexts of SNPs localized at both extremities of QTL CIs were blasted against reference genome GDDH13 v1.1 (using INRAE JBrowse tool) to retrieve QTL physical borders. Physical positions of virtual markers were estimated according to the physical positions of flanking SNP markers. The CI representing the shortest genetic region for each QTL was used to look for candidate genes involved in plant-pathogen interaction. For all QTLs, lists of genes along with their functional annotations and orthologs in *Arabidopsis thaliana* were extracted from GDDH13 v1.1 gene annotation (https://iris.angers.inra.fr/gddh13/the-apple-genome-downloads.html).

Transcriptomic datasets were retrieved from a previous experiment performed on offspring of the base population of the ‘TxF’ progeny ^29^. Four individuals for each of four classes were pooled according to the presence/absence of favorable alleles at three QTLs: qT1, qF11 and qF17. The four classes were constructed depending on the combination of QTLs present with either ‘NoScabQTL’, ‘qT1’, both ‘qF11qF17’ and all three ‘qT1qF11qF17’. This design allowed to test for the effects of QTLs on gene transcription before and after inoculation of *V. inaequalis* with the ‘EU-B04’ isolate (see Bénéjam et al. (2024) ^29^ for additional information).

For all candidate genes underlying the three QTLs qT1, qF11 and qF17, we thus retrieved fold changes and corrected p-values for genes showing differential transcription before and after *V. inaequalis* inoculation according to the genotypic classes. For instance, for genes within the qT1 range, we analyzed differential transcription between both qT1 and qT1qF11qF17 classes against the ‘NoScabQTL’ class, but also between qT1qF11qF17 class against the qF11qF17 class, at 0 dpi (before *V. inaequalis* inoculation) and 5 dpi (after *V. inaequalis* inoculation). Candidate genes with an absolute fold change (|FC|) > 1.5 and a Benjamini–Hochberg adjusted p-value (BH method) < 0.05 were selected for further analysis ^46^.

For the qT13 and qF3 QTLs, no transcriptomic data were available to study the expression of candidate genes contained in the CIs.

## Results

### 1. Broadening the mapping population and recombinant detection

Among the 1,970 genotyped seedlings, 1,101 displayed at least one recombination event within the five target QTL regions, thereby increasing mapping precision relative to Bénéjam et al. (2021) ^18^. Recombinant counts were notably higher for qT1 and qF11 (188 and 432 individuals, respectively; Table 2), substantially refining the genetic resolution.

**Table 2.** Summary of number of individuals presenting recombination events in confidence intervals of resistance quantitative trait loci.

|  | Recombinant number from<br>Bénéjam et al. (2021) |  | Recombinant number in this<br>study |  |
| --- | --- | --- | --- | --- |
| qT1 | 7 | 2% | 188 | 9% |
| qT13 | 16 | 6% | 168 | 8% |
| qF3 | 47 | 17% | 364 | 18% |
| qF11 | 43 | 16% | 432 | 21% |
| qF17 | 30 | 11% | 229 | 11% |
| Progeny size | 266 |  | 1,970 |  |

### 2. Phenotypic variation and symptom characterization

Phenotypes revealed a clear segregation pattern within the extended ‘TxF’ progeny. In the ‘Vi-B04’ experiment, out of the 1,970 seedlings, AUDPC values showed a distribution with a bimodal tendency (Fig 1A), with a large subset of ∼300 seedlings displaying near-zero values, indicative of strong resistance. The distribution of AUDPC showed a maximum value of 109 and a median of 44.6. When individuals carrying the favourable qT1 allele (n = 1,249) were removed, the remaining distribution became unimodal and continuous (Fig 1B). Median AUDPC value rose to 63.5, demonstrating the effect of qT1 face to this isolate. In contrast, adjusted AUDPC values obtained with ‘Vi-Z14’ experiments, induced a unimodal distribution across the 1,101 recombinants ranging from 0 to 112 AUDPC values, with a median of 55.8 (Fig 1C). The increase of median AUDPC in ‘Vi-Z14’ experiments illustrates the ability of the ‘09BCZ014’ to partially overcome the QTL qT1.

**Fig 1.**
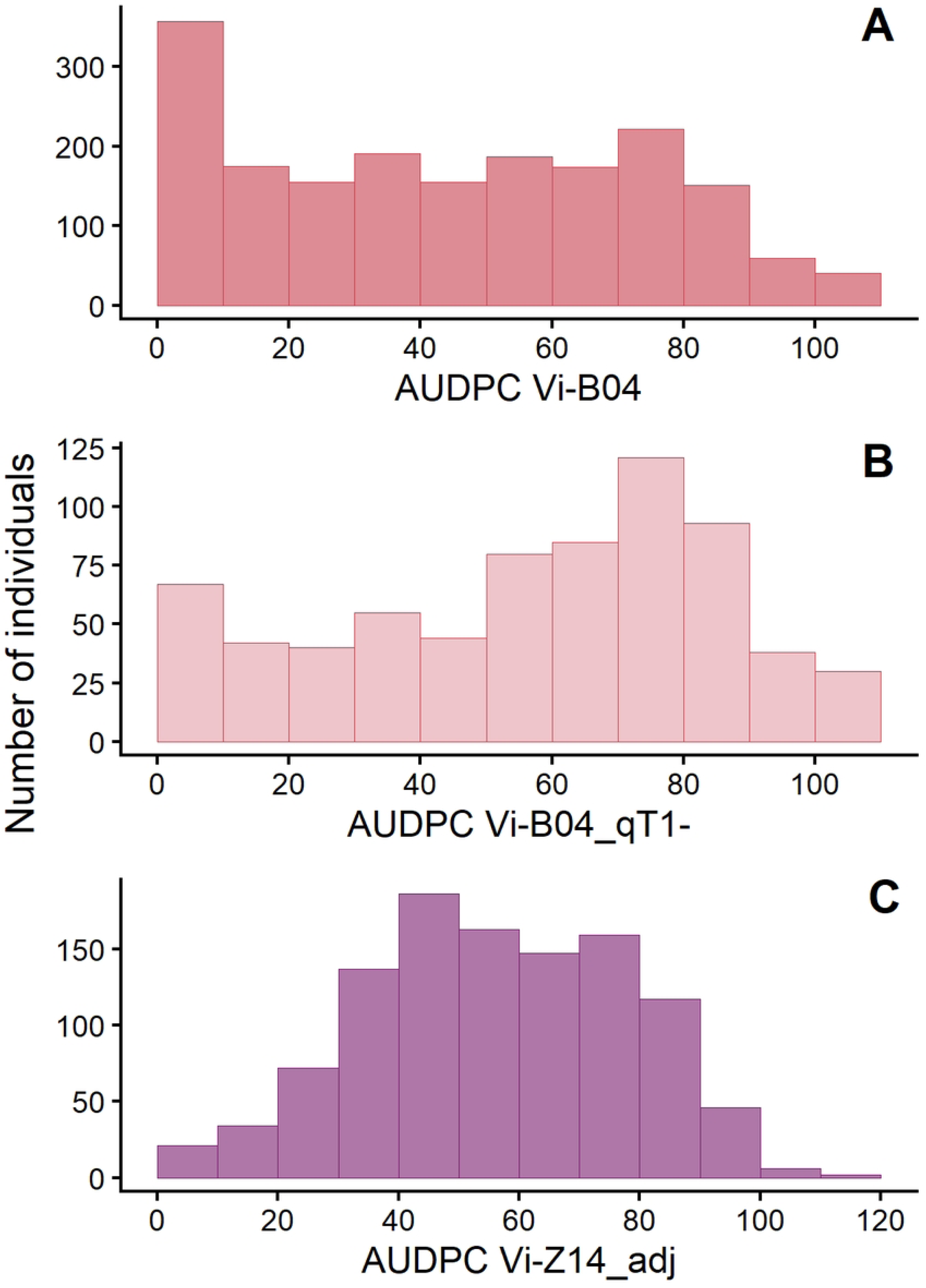
Distribution of susceptibility symptoms of the extended ‘TxF’ progeny. Adjusted Area Under the Disease Progress Curves (AUDPC) variation for the experiment with ‘EU-B04’ isolate (A) and experiment with the ‘09BCZ014’ isolate (C) are shown. Distribution of AUDPC estimated with ‘EU-B04’ isolate (B) is also represented in the subpopulation qT1- (i.e., individuals selected as not carrying the resistance allele of the major quantitative trait locus qT1).

Additionally, leaf resistance responses were scored in the ‘Vi-B04’ experiment through the assessment of chlorosis and crispation, each ranging from 0 to 7. Over 80% of genotypes exhibited both reactions (scores > 1). These two symptoms were strongly correlated (r = 0.649, p < 0.001) and inversely associated with disease severity (AUDPC; Fig 2). However, a substantial amount of genotypes was resistant (low AUDPC values) without presenting resistance symptoms, as illustrated by the low correlation scores between AUDPC and resistance phenotypes (r = −0.405 and −0.368).

**Fig 2.**
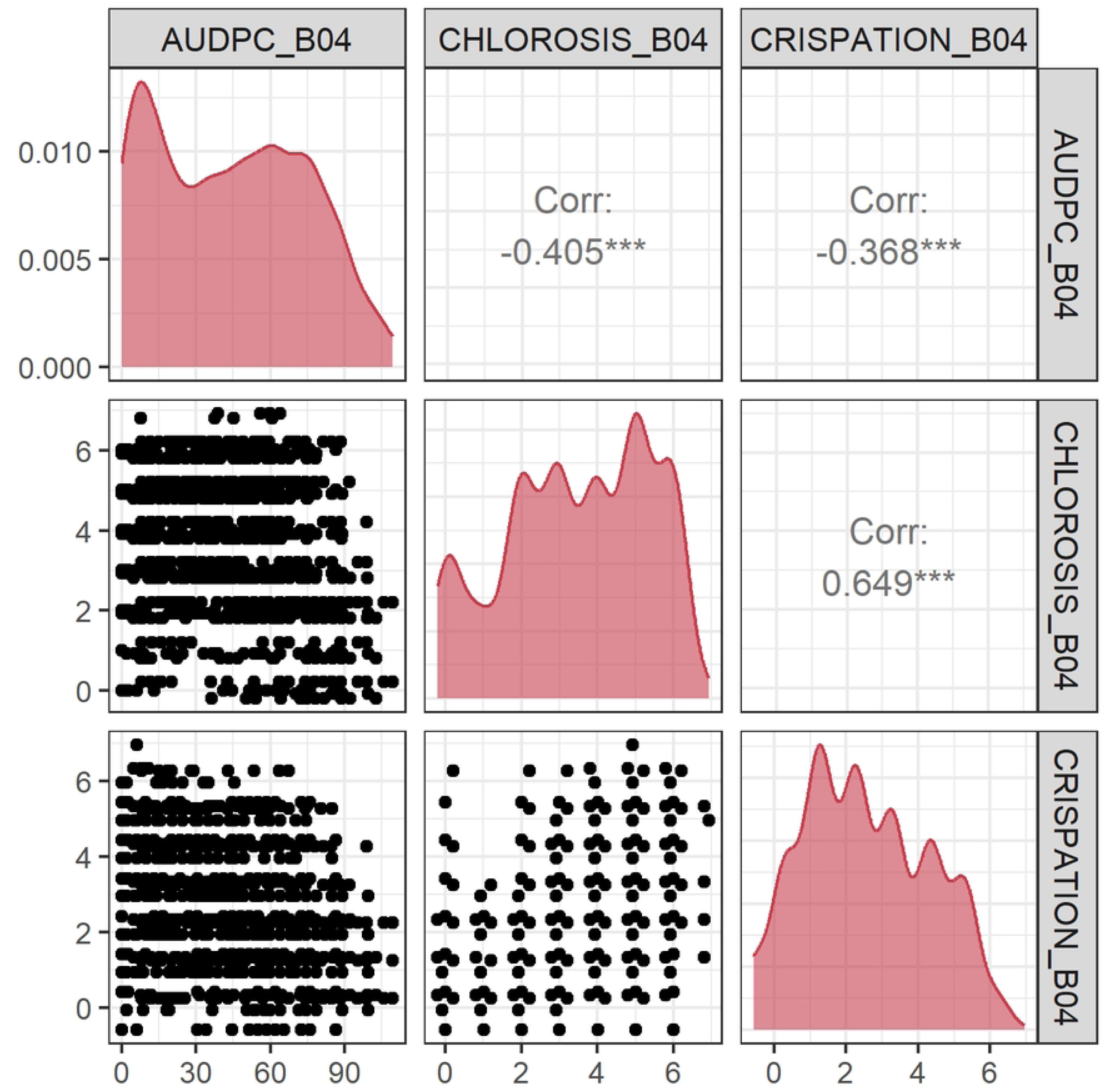
Relationship between resistance and susceptibility symptoms of the extended ‘TxF’ progeny after inoculation with isolate ‘EU-B04’. Pearson coefficient of correlation (Corr) are indicated for each couple of variables. Three variables measured in ‘Vi-B04’ experiment are presented: Area Under the Disease Progress Curve (AUDPC), chlorosis and crispation scoring.

### 3. QTL validation and fine mapping

QTL mapping analyses with ‘Vi-B04’ experiment variables were conducted on the entire progeny (1,970 seedlings) while only recombinants (1,101 seedlings) were considered for the analysis of AUDPC from ‘Vi-Z14’ experiment. Four of the five targeted QTLs were validated across experiments and phenotypic variables (qT1, qF11, qF17, and qT13) (Fig 3, S1 Table and S1 Fig). For the QTL qF3, crispation variable reached the detection threshold (LOD > 1.8) with a maximum value of 3.3, anchored by the virtual marker c3.loc15 (15 cM). Another signal was detected at the border of the studied interval (20 cM) with a value of 1.88 obtained with AUDPC from ‘Vi-Z14’ experiments. Despite dense marker coverage, this variable did not lead to the refinement of qF3 region and other phenotypic variables (chlorosis and AUDPC from ‘Vi-B04’ experiment) failed to reach the detection threshold (S1 Table and S1 Fig).

**Fig 3.**
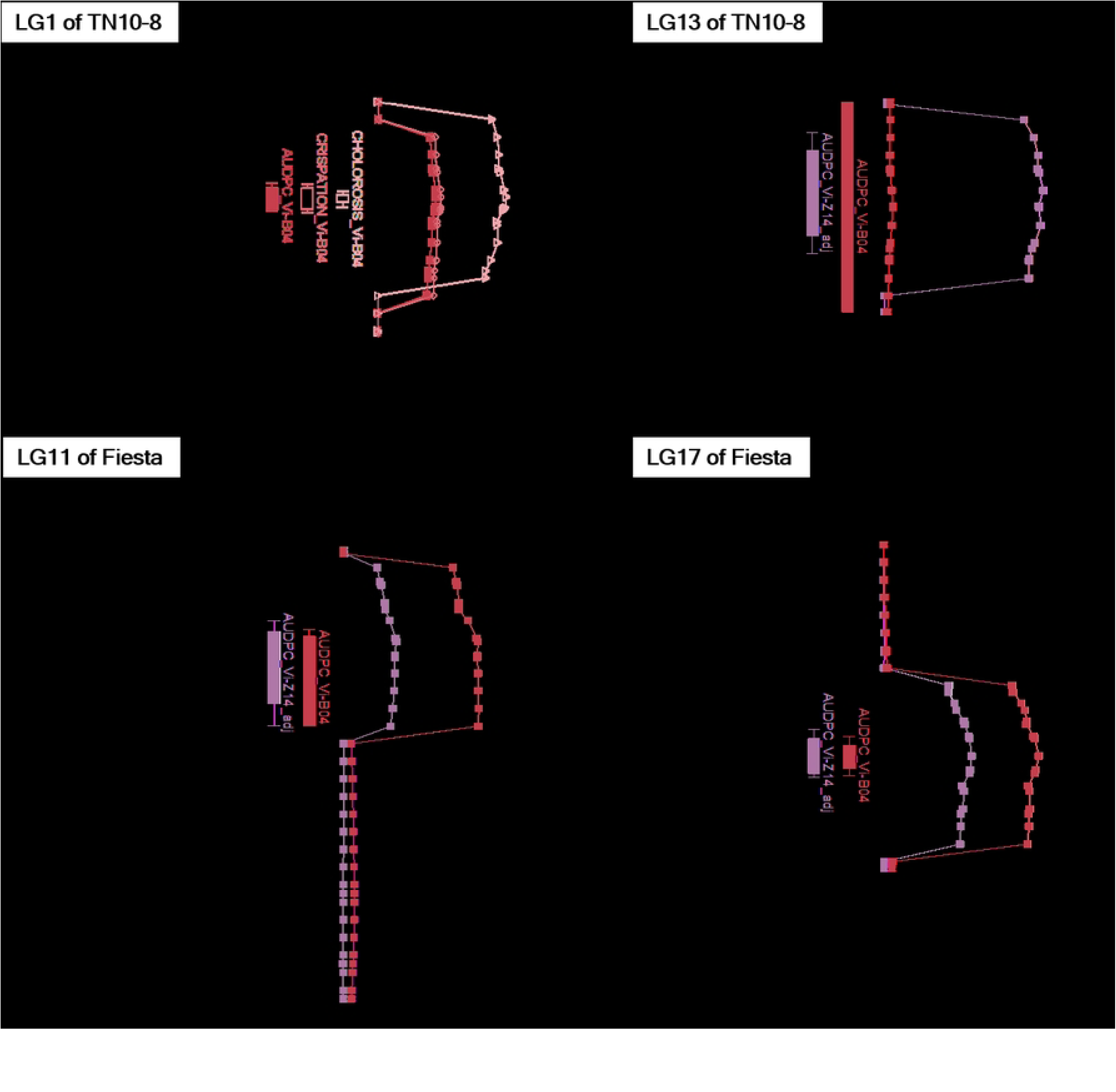
Genetic maps and LOD curves of the fine-mapped scab resistance quantitative trait loci of the extended ‘TxF’ progeny. Genetic position (in centiMorgans) and marker name used in this study are respectively indicated on the left and right of linkage groups (LG). Marker name associated to maximum LOD value for each QTL are bolded. 2-LOD and 1-LOD support QTL confidence interval are represented by vertical lines and solid rectangles, respectively.

The QTL qT1 was detected in ‘Vi-B04’ experiment for all phenotypic traits, displayed the highest significance (LOD = 202) when using chlorosis as the trait. Its 2-LOD CI was narrowed to 1.1 cM (∼600 kb on the GDDH13 genome), anchored by marker AX-115385376. Similar intervals were estimated with AUDPC and crispation variables from this experiment.

For qF11 and qF17, the shortest 2-LOD CIs were obtained with AUDPC from ‘Vi-Z14’ experiments. The qF11 region was delimited to a 6 cM region (∼2.5 Mb), while qF17 was refined to 3.5 cM (∼1.2 Mb). The most probable position of these QTLs (i.e. the positions with highest LOD scores) were anchored by markers AX-115185079 and AX-115185087 for qF11 (5.2 cM), and by the virtual marker c17.loc12 (12 cM) for qF17. These two QTLs were also identified in the ‘Vi-B04’ experiment using AUDPC and exhibited longer intervals.

Finally, qT13 was detected exclusively under ‘Vi-Z14’ inoculation, within a 7 cM (∼2 Mb) region on chromosome 13, anchored by the AX-115187347 marker. Although it did not reach the LOD threshold, the AUDPC from the ‘Vi-B04’ experiment yielded a LOD score of 1.78 on this marker.

### 4. Epistatic interaction between qF11 and qF17

Consistent with prior findings, seedlings carrying favorable alleles at both qF11 and qF17 exhibited significantly lower AUDPC values than all other genotypic classes (p < 0.001; S2 Fig). Under ‘Vi-B04’, median AUDPC dropped from 50–60 in single-allele carriers to ∼15 in double carriers; under ‘Vi-Z14’, values decreased from ∼60 to ∼30. Also, single-allele exhibited susceptibility levels similar to those of non-carriers. These patterns confirm a synergistic effect between qF11 and qF17.

### 5. Identification of candidate genes and defense mechanisms

To pinpoint plausible resistance determinants, we focused on genes located within the refined CIs and showing differential expression in relevant genotypic classes (before and five days after inoculation).

#### qT1 region

The narrow 590 kb qT1 interval encompassed 89 annotated genes, 10 of which were differentially expressed (Table 2). Most of these were upregulated following inoculation in resistant (qT1+) genotypes. Notably, five clustered genes (*MD01G1178700, MD01G1179300, MD01G1179700, MD01G1179800,* and *MD01G1180500*) encode Leucine-Rich Repeat Receptor-Like Proteins (LRR-RLPs), classical sensors of fungal or oomycete invasion. This transcriptional pattern supports the hypothesis that *qT1* could represent either a functional variant or a paralog of *Rvi6* with partial quantitative activity.

**Table 2.** Annotation and relative expression of genes underlying the QTL qT1 confidence interval before (0 dpi) or five days (5 dpi) after infection with *Vi* in different genetic background of pooled-samples from the extended ‘TxF’ progeny.

| QTL | Gene ID | Annotation<br>Swissprot | TAIR Name | Biological process | 0 dpi |  |  | 5 dpi |  |  | 0 dpi versus 5 dpi |  |  |  |
| --- | --- | --- | --- | --- | --- | --- | --- | --- | --- | --- | --- | --- | --- | --- |
|  |  |  |  |  | NoScabQTL/<br>qT1 | NoScabQTL/<br>qT1qF11qF17 | qF11qF17/<br>qT1qF11qF17 | NoScabQTL/<br>qT1 | NoScabQTL/<br>qT1qF11qF17 | qF11qF17/<br>qT1qF11qF17 | NoScabQTL | qT1 | qF11qF17 | qT1qF11qF17 |
| qT1 | MD01G1173000 | Phytosulfokine<br>receptor 1 | AT1G74190.1 | Signal transduction | 2,4 | - | - | - | - | - | - | 2,2 | - | - |
|  | MD01G1177000 | Ethylene-<br>responsive<br>transcription<br>factor 2 | AT4G17500.1 | Cellular process /<br>response to nematode | - | - | - | - | - | - | - | 1,6 | - | - |
|  | MD01G1177100 | Ethylene-<br>responsive<br>transcription<br>factor 5 | AT5G51190.1 | Response to heat | - | - | - | 1,67 | - | - | - | - | - | - |
|  | MD01G1178700 | LRR receptor-like<br>serine/threonine<br>-protein kinase<br>FLS2 | AT2G34930.1 | Defense response to<br>fungus and oomycetes | - | - | - | - | - | - | - | 2,7 | - | 2,6 |
|  | MD01G1178900 | Receptor-like<br>protein 12 | AT1G07390.3 | Signal transduction | - | - | - | - | - | - | - | 2,7 | - | - |
|  | MD01G1179300 | LRR receptor-like<br>serine/threonine<br>-protein kinase<br>GSO2 | AT2G34930.1 | Defense response to<br>fungus and oomycetes | - | - | - | - | - | - | 1,7 | 1,5 | - | - |
|  | MD01G1179700 | Probable LRR<br>receptor-like<br>serine/threonine<br>-protein kinase<br>IRK | AT2G34930.1 | Defense response to<br>fungus and oomycetes | - | - | - | - | - | - | - | 2,8 | - | 5,4 |
|  | MD01G1179800 | Receptor-like<br>protein 12 | AT2G34930.1 | Defense response to<br>fungus and oomycetes | - | - | - | - | - | - | - | 2,3 | - | - |
|  | MD01G1180500 | Receptor-like<br>protein 12 | AT2G34930.1 | Defense response to<br>fungus and oomycetes | - | - | -1,5 | - | - | - | - | 2,3 | - | 2,4 |
|  | MD01G1181100 | Probable inactive<br>leucine-rich<br>repeat receptor<br>kinase XIAO | AT2G34930.1 | Defense response to<br>fungus and oomycetes | - | 2,3 | - | 3,19 | 3 | 3,3 | - | - | - | - |

#### qF11 and qF17 regions

The refined qF11 (2.6 Mb) and qF17 (1 Mb) intervals contained 304 and 130 genes, respectively. Fourteen were differentially expressed in each region (Table 3). In qF11 interval, several DEGs (*MD11G1025200–MD11G1033300*) were constitutively upregulated in resistant genotypes at both time points, suggesting baseline activation of signaling pathways rather than induced defense. Their functions, which are linked to RNA interference and signal transduction, suggest the existence of a regulatory layer that modulates defense gene networks by inducing kinases, ligases, and helicases.

**Table 3.** Annotation and relative expression of genes underlying the QTL qF11 and qF17 confidence intervals before (0 dpi) or five days (5 dpi) infection with *Vi* in different genetic background of pooled-samples from the extended ‘TxF’ progeny.

| QTL | Gene ID | Annotation<br>Swissprot | TAIR Name | Biological process | 0 dpi |  |  | 5 dpi |  |  | 0 dpi versus 5 dpi |  |  |  |
| --- | --- | --- | --- | --- | --- | --- | --- | --- | --- | --- | --- | --- | --- | --- |
|  |  |  |  |  | NoScabQTL/<br>qF11qF17 | NoScabQTL/<br>qT1qF11qF17 | qT1/<br>qT1qF11qF17 | NoScabQTL/<br>qF11qF17 | NoScabQTL/<br>qT1qF11qF17 | qT1/<br>qT1qF11qF17 | NoScabQTL | qT1 | qF11qF17 | qT1qF11qF17 |
| qF11 | MD11G1022100 | - | - | - | 2,4 | 2,2 | - | - | - | - | - | - | - | - |
|  | MD11G1025200 | G-type lectin S-receptor-like serine/threonine-protein kinase LECRK3 | AT5G60900.1 | - | 3,4 | 3,3 | 2,2 | 3,4 | 4,2 | 2,1 | - | - | - | - |
|  | MD11G1025300 | DEAD-box ATP-dependent RNA helicase 20 | AT1G55150.1 | RNAi-mediated immune response | 8,7 | 8,7 | 8,8 | 8,5 | 8,5 | 7,7 | - | - | - | - |
|  | MD11G1025500 | G-type lectin S-receptor-like serine/threonine-protein kinase LECRK3 | AT5G60900.1 | - | 2,7 | 2,4 | - | 2,4 | 3,3 | - | - | 1,8 | - | 1,5 |
|  | MD11G1026100 | DEAD-box ATP-dependent RNA helicase 20 | AT1G55150.1 | RNAi-mediated immune response | 10,3 | 10,2 | 10,4 | 10,8 | 10,9 | 12,4 | - | - | - | - |
|  | MD11G1027500 | Beta-glucosidase 12 | AT2G44480.1 | Carbohydrates metabolic process | - | - | - | - | - | - | 1,8 | 2,5 | - | - |
|  | MD11G1027800 | TIR domain-containing adapter molecule 1 | AT4G12070.1 | - | - | - | - | - | - | - | - | 2,2 | - | 2,7 |
|  | MD11G1031100 | F-box protein CPR30 | AT4G12560.1 | Regulation of defense response | - | - | - | - | 2,6 | - | - | - | - | - |
|  | MD11G1031200 | #N/A | #N/A | - | - | - | - | -2,0 | - | - | - | - | - | - |
|  | MD11G1033300 | Serine/threonine-protein kinase STY17 | AT3G22750.1 | Signal transduction | 6,0 | 6,0 | 3,4 | 5,3 | 4,5 | 2,9 | - | - | - | - |
|  | MD11G1038900 | Fructose-bisphosphate aldolase 5, cytosolic | AT4G26530.1 | Carbohydrates metabolic process | - | - | - | -1,6 | - | - | - | - | - | - |
|  | MD11G1039300 | Heavy metal-associated isoprenylated plant protein 16 | AT3G07600.1 | - | - | -3,5 | -3,8 | - | - | - | - | - | - | - |
|  | **MD11G1043900** | C6 finger domain transcription factor tcpZ | AT1G07745.1 | Cellular process: DNA repair | - | - | -3,1 | - | - | - | - | - | - | - |
|  | **MD11G1049600** | Cytochrome P450 714C2 | AT5G24910.1 | - | - | - | - | - | - | - | - | - | - | 2,8 |
| **qF17** | **MD17G1098400** | Trophinin | AT5G07570.1 | - | 1,5 | - | - | - | - | - | - | - | - | - |
|  | **MD17G1099000** | WRKY transcription factor 42 | AT4G04450.1 | Leaf senescence regulation | - | - | - | - | - | - | - | - | - | 1,7 |
|  | **MD17G1099500** | Cellulose synthase-like protein B3 | AT2G32530.1 | Cellulose biosynthetic process | - | - | - | - | - | - | - | 2,0 | - | - |
|  | **MD17G1099600** | Cellulose synthase-like protein H1 | AT2G32530.1 | Cellulose biosynthetic process | - | - | - | - | - | - | - | 1,7 | - | - |
|  | **MD17G1100100** | Fatty acid oxidation complex subunit alpha | AT4G20720.1 | - | - | 1,6 | - | - | - | - | - | - | - | - |
|  | **MD17G1101300** | Condensin complex subunit 2 | AT2G32590.1 | Mitotic chromosome condensation | - | 2,1 | 3,7 | - | - | - | - | - | - | - |
|  | **MD17G1101700** | UPF0753 protein YbcC | AT1G77130.1 | Plant secondary cell wall biogenesis | - | - | - | - | - | - | - | 1,8 | - | - |
|  | **MD17G1102000** | Ribonuclease 3-like protein 3 | AT4G15417.1 | RNA processing | - | - | - | - | - | - | - | 2,5 | - | - |
|  | **MD17G1104200** | UDP-glucose:glycoprotein glucosyltransferase | AT5G04700.1 | - | -12,2 | -12,0 | -11,8 | -5,9 | -8,4 | -8,5 | - | - | - | - |
|  | **MD17G1104700** | DEAD-box ATP-dependent RNA helicase 45 | AT3G09620.1 | mRNA splicing | - | - | - | - | - | -5,0 | - | - | - | - |
|  | **MD17G1105000** | Uncharacterized protein DDB_G0275933 | AT5G16060.1 | - | -2,0 | -2,1 | -1,8 | -2,1 | -2,2 | -2,2 | - | - | - | - |
|  | **MD17G1105800** | Palmitoyl-monogalactosyl diacylglycerol delta-7 desaturase, chloroplastic | AT3G15850.1 | Fatty acid biosynthetic process | - | - | - | - | - | - | - | - | - | 1,6 |
|  | **MD17G1106300** | 1-aminocyclopropane-1-carboxylate oxidase | AT1G05010.1 | Ethylene biosynthetic process / response to fungus | - | - | - | - | - | - | - | 2,0 | - | 2,1 |
|  | **MD17G1109600** | Chlorophyll a-b binding protein 40, chloroplastic | AT1G29930.1 | Photosynthesis / response to light stimulus | - | - | 4,3 | - | - | - | - | 3,6 | - | - |

Conversely, two DEGs in qF17 interval (*MD17G110420, MD17G1105000*) were constitutively downregulated in resistant individuals. Although their annotations remain uncertain (one glucosyltransferase and one uncharacterized protein), predicted enzymatic roles could cover various functions. Interestingly, several DEGs in both regions were induced by inoculation, but none of them was specific to the qF11qF17 genotypic class.

#### qT13 region

qT13 spanned a 2 Mb interval containing 289 genes, 22 of which lacked functional annotation (S2 Table).

## Discussion

The broadening of the biparental population and the analysis of nearly 2,000 seedlings allowed the refinement and validation of several resistance QTLs previously reported in the ‘TxF’ progeny. Among the five QTLs studied, qT1, qF11, qF17, and qT13 were validated under at least one experimental condition, while qF3 could not be confirmed despite an increased number of individuals. The ‘Vi-B04’ experiment revealed a somewhat bimodal distribution of resistance phenotypes mainly driven by the presence of the qT1 QTL. The joint analysis of differential gene expression within refined CIs identified several candidate genes, including RLPs in qT1 and constitutively expressed genes in qF11 and qF17. Epistatic interaction between qF11 and qF17 was confirmed, supporting the functional association of their underlying genes in resistance expression.

### Experimental validation limits and interpretation of QTL stability

The absence of qF3 detection and the variability observed in QTL expression across experiments underline the context-dependence of QTL detection. For qF3, QTL detection was successful only for the crispation variable, whereas no QTL were identified for chlorosis, in contrast to other detections (Fig 3) and despite their correlation (Fig 2). In addition, given the low LOD score, this result should be interpreted with caution and confirmed in independent experiments with crispation variable. Although the overall results are mixed, we observed an increase in LOD values towards the end of the targeted genomic region, which was even more pronounced when using AUDPC from the ‘Vi-Z14’ experiment (S1 Fig). This pattern may suggest that the QTL is located slightly downstream of the initially targeted region. Such an interpretation would be consistent with previous findings, in which the position of the highest LOD scores varied among QTL detections performed with two different *Vi* isolates ^18^. Although,

Bénéjam et al. (2021) ^18^ studied 267 individuals, out of them 17% presented recombination events in qF3 interval, which is similar to our study (Table 2). However, without biological replicates, it is possible that phenotypic responses were less precise in this study, leading to an absence of detection of low effect QTLs, such as qF3. Another possible explanation is that qF3 may have been a false positive in the previous study, in which it was detected for the first time ^18^. Further analyses using a higher number of markers and replicates would be required to clarify this point.

The non-validation of qT13 in ‘Vi-B04’ experiment contrasts with its previous identification by Bénéjam et al. (2021) ^18^ in both isolate contexts. However, the improved precision of its mapping interval supports the robustness of the locus itself. In addition, QTL detection with AUDPC from ‘Vi-B04’ experiment almost reach the LOD threshold (Fig 3). Increasing the level of replication in future experiments, notably with the isolate ‘EU-B04’ could help refine the localization of the QTL by estimating more reliable AUDPC values. The current detection of qT13 only in ‘Vi-Z14’ experiments may also indicate isolate-specific activation or environmental modulation of gene expression. Despite the reduction of the confidence interval, 283 genes still fall within the boundaries of qT13 (S2 Table). With only seven markers on this QTL in this study, it would be relevant to further increase marker density in this region, particularly around the marker AX-115187347 (5.89 cM), since the peak of LOD score was located near it (S1 Table). Moreover, no transcriptomic data are available for this QTL, preventing the initiation of gene number reduction and the investigation of potential gene functions. Due to its efficacy against at least isolate ‘09BCZ014’, ‘TN10-8’ remains an attractive parent for breeding.

In addition, several non-genetic factors may have contributed to this instability. Differences in environmental conditions between experiments, and particularly the use of rootstocks for individual replication in earlier studies, could have modified physiological parameters and biased disease symptom expression. Ontogenic effects and developmental stage of seedlings at inoculation are known to strongly influence the expression of resistance loci, as reported in apple and other perennial hosts ^17,47,48^. Here, scab inoculations were carried out sequentially on the same plants, which were pruned between each inoculation. Similarly, Soufflet-Freslon et al. (2008) ^17^ reported differences in AUDPC distributions of a progeny when screened successively in two scab experiments using the same *Vi* isolate, highlighting differential resistance expression depending on the physiological state of the plants. Collectively, these results suggest that the variability in QTL validation partly reflects the ontogenic and polygenic nature of apple scab resistance.

### Functional interpretation and convergence of molecular mechanisms

The co-localization of differentially expressed genes (DEGs) with refined QTL intervals can reveal potential functions of these loci. The working hypothesis underlying this analysis is that the QTL effect results from allele-specific differences in gene expression, caused by sequence variation in regulatory regions of the underlying gene. Under this model, differential transcription between alleles would drive the observed phenotypic variation. Alternative mechanisms cannot be excluded, including coding sequence variation leading to structural differences in the encoded protein without expression differences ^49^, or epigenetic regulation generating allele-specific expression in the absence of sequence polymorphism ^50^. In this context, RNA-seq data provide a relevant framework to investigate the expression-based hypothesis and to prioritize genes within the QTL CI, notably by excluding genes not expressed under our experimental conditions.

For qT1, due to its specificity, large-effect and physical localization on the genome, it has already been hypothesized to be an allelic variant or a paralog of the major/R gene *Rvi6* ^16,18^. Moreover, qT1 often leads to a hypersensitive response ^20^ and transcriptomic datasets highlighted a large amount of DEGs (∼1,500) when comparing genotypes carrying susceptibility and resistance alleles of qT1 infected with ‘EU-B04’ isolate ^29^. Here, we refined the CI of qT1 to a ∼600 kb region (Fig 3, S1 Table) and analyzed DEGs within this interval. More precisely, several RLPs were upregulated upon *Vi* inoculation (Table 2), a pattern consistent with canonical R-gene–mediated perception ^51^. This supports the hypothesis that qT1 might be an allele or a paralog of the well-known *Rvi6* locus, with different specificities ^52^. A comparative genomic analysis of the potential allelic series in *Rvi6* region, comprising *Rvi6* cloned in ‘Florina’ ^8,9,53^ and also present in recently sequenced ‘Prima’ and ‘Priscilla’ ^54^, *Rvi17* in ‘Antonovka’ ^55^, *Vhc1* in ‘Honeycrisp’ ^56^ and qT1 in ‘TN 10-8’, would help confirm this co-localization and clarify their possible redundancy or evolutionary divergence. A definitive test would examine segregation in progeny from crosses between parents carrying different putative alleles of these R genes/QTLs. Detection of both “alleles” in offspring would indicate recombination, showing they are distinct loci rather than alternative alleles. In a perennial species, such crosses are labor- and time-intensive, and testing all pairwise allele combinations is practically difficult.

For qF11 and qF17, CIs were narrowed to 2.6 Mb and 1 Mb (Fig 3, S1 Table), respectively. Their epistatic interaction was also confirmed (S2 Fig), consistent with other studies ^18–20^. Integration of RNA-seq data provided additional insight into the biological mechanisms potentially underlying the interaction between these QTLs, possibly through coordinated transcriptional regulation. Within the qF11 interval, genes putatively involved in RNAi signaling were expressed, whereas the qF17 interval showed repression of genes with currently unknown functions (Table 3). In both regions, expression differences were primarily associated with genotypic classes rather than with *Vi* inoculation, supporting the hypothesis that these QTLs are constitutively expressed and may contribute to basal resistance ^20^. Nevertheless, improved gene annotation will be necessary to better characterize the molecular basis linking these two loci. The use of haplotype-resolved parental genomes would provide an ideal framework for identifying genes underlying QTLs. The causal gene may be absent from one haplotype in the case of hemizygous genes, or even missing from the reference genome if the sequenced genotype does not carry the QTL allele.

## Conclusions

The development of new markers and the screening of nearly 2,000 seedlings helped consolidate our understanding of the genetic architecture of scab resistance in the ‘TxF’ progeny, validating four QTLs with a promising breadth of action. The improved precision of QTL intervals, comprised between 2.6 Mb and 600 kb, and integration of transcriptomic data strengthen the genetic framework for downstream applications. Future efforts should focus on (i) confirming the co-localization between qT1 and *Rvi6*, (ii) functionally characterizing qT13, and (iii) elucidating the biological basis of the epistatic interaction between qF11 and qF17 integrating other phenotypic data. In breeding terms, the combination of these loci through marker-assisted selection, supported by the development of new SNP markers, represents a promising path toward durable resistance in apple.

## Author contributions

Romane Lapous: Investigation, Data Curation, Validation and Visualization of results, Writing – Original Draft Preparation, Review & Editing. Camille Haquet: Investigation, Data Curation, Formal analysis, Writing – Review & Editing. Caroline Denancé: Investigation, Data Curation. Juliette Bénéjam: Validation and Visualization of results, Writing – Review & Editing. Laure Perchepied: Validation and Visualization of results, Writing – Review & Editing. Kaat Hellyn: Resources. Hélène Muranty: Conceptualization, Funding Acquisition, Writing – Review & Editing. Charles-Eric Durel: Conceptualization, Funding Acquisition, Writing – Review & Editing. Julie Ferreira de Carvalho: Investigation, Conceptualization, Funding Acquisition, Project Administration, Supervision, Writing – Original Draft Preparation, Review & Editing.

## Acknowledgements

This work was supported by (i) a grant to the METAdiVERSE project from the BAP (Plant biology and breeding) division of INRAE and by (ii) a grant from the French government managed by the Agence Nationale de la Recherche (ANR) as part of the Programme Prioritaire de Recherche “Cultiver et Protéger Autrement” under the reference ANR-20-PCPA-0003 (CapZeroPhyto project). Romane Lapous was supported by a Ph.D. fellowship from the BAP division of INRAE and the ‘Pays de la Loire’ region (France). Camille Haquet was supported by a GIS Fruit grant. The authors greatly thank the PHENOTIC platform (https://doi.org/10.17180/YKBZ-2V85) for carefully taking care of the plant material. The authors would like to thank the Biological Resource Center “RosePom - Pome Fruits and Roses” (https://eng-irhs.angers-nantes.hub.inrae.fr/shared-facilities/genetic-resources/crb-pome-fruit-and-rose) and associated staff for maintaining the plant material and associated datasets used in the present article. We are also grateful to the Horticole Experimental Unit (https://doi.org/10.15454/1.5573931618268674E12) for maintaining the young trees derived from seedlings. For generating the seeds used in this study, we thank present and former staff of the VaDiPom team (IRHS). We thank Dr. Diego Micheletti for sharing the alignment of sequencing data to the GGDH13 genome. For selecting and developing SNP markers, we thank Aurélien Petiteau and the ANAN platform (SFR QuaSaV, Univ Angers). Finally, the authors wish to thank M. Tiret for thorough reviewing of the original draft as well as all the people who helped in the sampling of leaf material and apple scab symptom scoring: L. Vitteaut, A. Daligault, A. Petiteau, B. Petit, R. Leclair, F. Lebreton.

## Supporting information

**S1 Fig.**
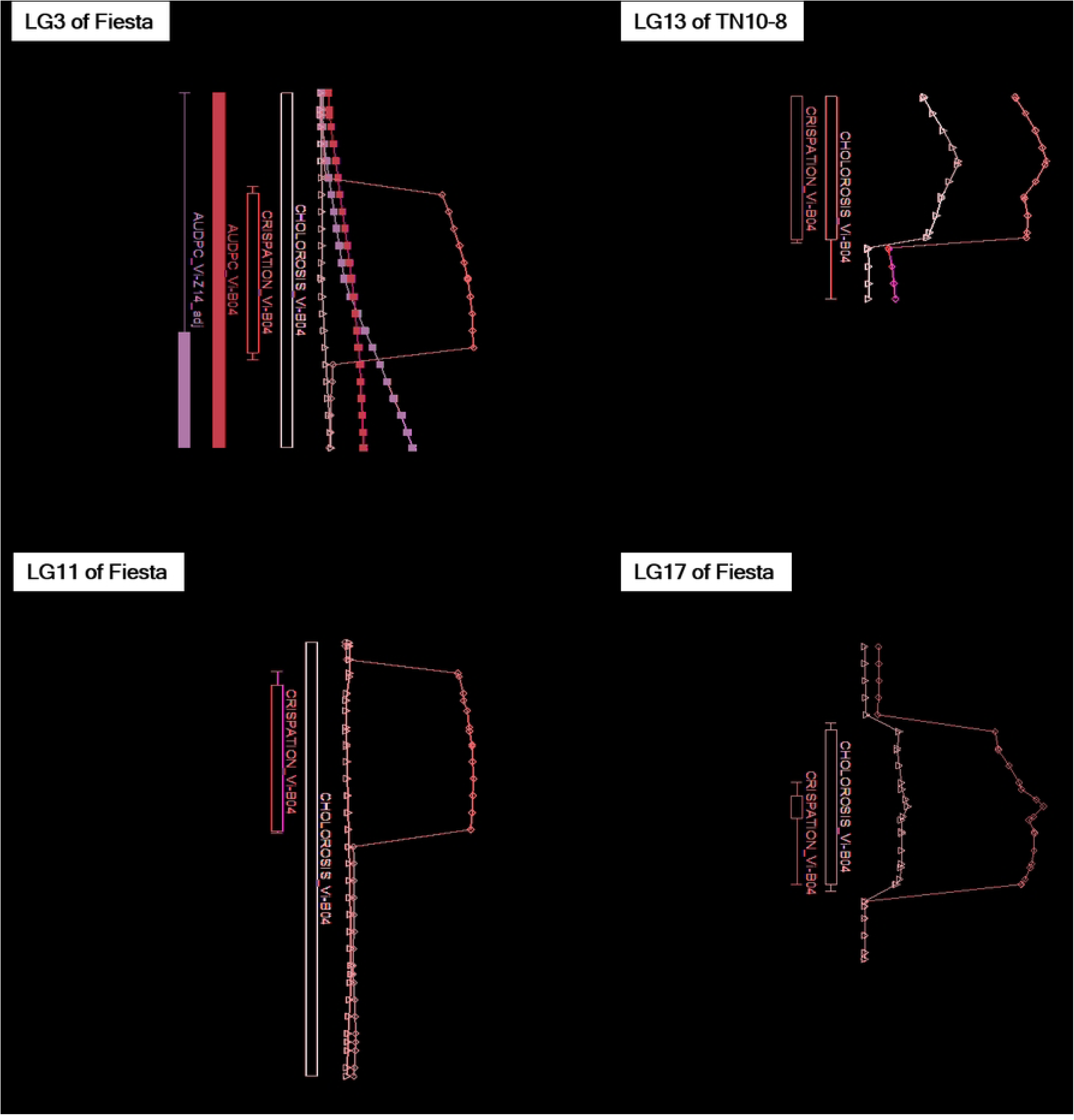
Genetic maps and LOD curves of minor variables used for scab resistance quantitative trait loci mapping of the extended TxF progeny. Genetic position (in centiMorgans) and marker name used in this study are respectively indicated on the left and right of linkage groups (LG). Marker name associated to maximum LOD value for each QTL are bolded. 2-LOD and 1-LOD support QTL confidence interval are represented by vertical lines and solid rectangles, respectively.

**S2 Fig.**
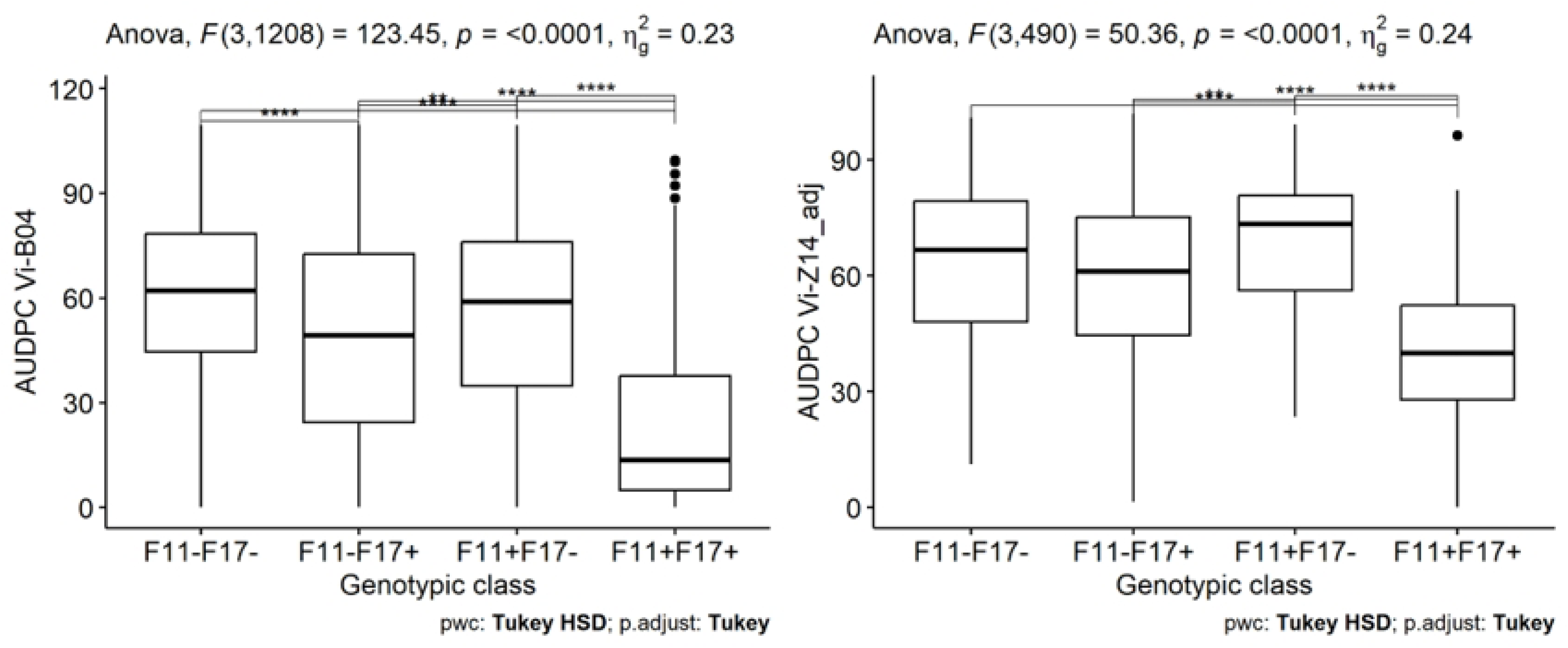
Distribution of AUDPC scores after scab infection with two isolates according to favorable allele of two epistatic QTL segregating in the extended TxF progeny. Genotypic classes are attributed according to allelic variant of marker associated to maximum LOD score of each QTL. Two variables are presented (adjusted AUDPC from ‘Vi-B04’ and ‘Vi-Z14’ experiments). Significant differences between classes have been evaluated through ANOVA test.

**S1 Table. Parameters associated with the fine-mapped quantitative trait loci (QTL) identified for scab resistance of the extended TxF progeny.** Virtual markers, for which physical positions were estimated, are also included (e.g., ‘c11.loc9’, which is placed on chromosome 11 at 9 centiMorgans). | Chr, chromosome, LOD, Logarithm of the odds; R², variance; CI, confidence interval.

**S2 Table. Gene list included in the refined confidence interval of qT13.** Gene names (‘Gene ID’) are extracted from the GDDH13 reference genome. | QTL, Quantitative Trait Loci.

**S1 File. Markers comprising the genetic map of the extended TxF progeny.** Virtual markers, for which physical positions were estimated, are included (e.g., ‘c11.loc9’, which is placed on chromosome 11 at 9 centiMorgans). Physical positions are extracted from the GDDH13 reference genome.

**S2 File. Phenotyping data used for QTL mapping in the extended TxF progeny.** Data were acquired across three experiments. One includes the inoculation with the *Vi* isolate ‘EU-B04’ and three phenotypic variables were scored. The two others experiments were conducted with the inoculation of isolate ‘09BCZ14’ and only sporulation symptoms were measured. Missing values are indicated with an asterisk (*).

**S3 File. Genotyping data used for QTL mapping in the extended TxF progeny.** For each SNP, alleles are coded as 1 or 2. Virtual markers, further imputed in genetic analyses, are not included. Missing values are indicated with an asterisk (*).

